# CortexDecoder: Efficient fMRI-to-Image Reconstruction from Cortical Flatmaps

**DOI:** 10.64898/2026.10.09.757766

**Authors:** Teo Imoto-Tar, Maxwell Fung, Arnav Gurha

## Abstract

Recent brain decoding work has shown the potential of Vision Transformers (ViTs) to learn mappings from neural representations to visual features. Here we present CortexDecoder, a framework that reconstructs viewed images from functional magnetic resonance imaging (fMRI) flatmaps, which are neural responses projected onto the cortical surface and unfolded into 2D images. We show that a frozen, pretrained cortical foundation model is sufficient for competitive reconstruction. A frozen CortexMAE encoder turns flatmap patches into brain tokens, and two lightweight cross-attention bridges map these tokens to semantic CLIP embeddings and low-level image features. The final image is generated using a frozen SDXL-unCLIP model which takes a low-level image reconstruction as an initial image and a predicted CLIP embedding used to condition the image generation. On Natural Scenes Dataset (NSD) Subject 1 reconstruction benchmarks, CortexDecoder is competitive with prior methods and the best for cortical flatmap methods without multi-subject pretraining. CortexDecoder achieves this performance while training far fewer parameters compared to prior methods.

## I. Introduction

Reconstructing visual experiences from non-invasive functional magnetic resonance imaging (fMRI) remains a central challenge in computational neuroscience. It also has applications in Brain-Computer Interfaces (BCIs), such as communication aids. With the availability of large, open-source fMRI-image datasets like the Natural Scenes Dataset (NSD) [2], fMRI has become a natural choice for studying visual decoding. The NSD provides paired fMRI responses together with natural images viewed by subjects, with repeated presentations of visual stimuli across trials. Each sample contains a cortical response corresponding to a viewed image, providing both neural and visual representations for learning fMRI-to-image mappings.

Our central question is whether we can efficiently decode visual stimuli from 2D cortical flatmaps. Most brain decoding frameworks follow one of two practices. Some approaches regress directly from raw 3D functional volumes, but this is challenging due to their high dimensionality, noisy signal, and computational cost. Alternatively, conventional approaches that flatten voxel responses into feature vectors or average responses over anatomical parcellations do not explicitly preserve spatial information. This motivated us to explore 2D flatmaps, which preserve cortical geometry while also being directly compatible with modern vision architectures.

We introduce CortexDecoder, a framework that decodes 2D cortical flatmaps into image stimuli using vision transformer (ViT) [3] models. Since flatmaps are inherently 2D images, we can partition neural activity into discrete patches. We use a frozen pretrained CortexMAE [4] encoder—an fMRI foundation model developed by MedARC which uses a masked autoencoder framework trained on cortical flatmaps. This encoder provides us with spatially organized neural representations. By utilizing the full sequence of spatially organized patch tokens from CortexMAE, our architecture preserves local cortical geometry and avoids collapsing the brain representation into a rigid 1D feature vector.

Similar to DETR (DEtection TRansformer) [5] and Brain-IT [6], our cross-attention bridge uses learned query vectors to map neural representations to visual feature spaces. Our cross-attention bridge is split into two modules to separately handle predicting semantic and structural visual information. The semantic branch predicts unpooled image CLIP [7] embeddings, which capture the concepts in the scene, whereas the low-level branch targets early visual features such as basic shapes and colors. Many decoding models struggle with low-level visual fidelity, often generating images that match semantically but are unfaithful in terms of pixel correlation and structural similarity. Incorporating a low-level branch targets this gap by providing structural guidance. Finally, we employ MindEye2’s [8] frozen SDXL-unCLIP model [9] to condition on both the low-level reconstruction and a semantic embedding to produce the final image.

We propose CortexDecoder, a reconstruction framework for cortical flatmap decoding which reads the full sequence of flatmap patch tokens with learned-query cross-attention. We also show that two trained lightweight bridges together with pretrained frozen foundation models are sufficient to achieve performance competitive with other fMRI-to-image decoding models.

## II. Related work

### A. fMRI-to-Image Reconstruction

Recent methods for fMRI-to-image reconstruction use fMRI data to predict the output embeddings of pretrained vision encoders such as CLIP. These output embeddings are then used to generate images with pretrained diffusion models. Notably, MindEye [11] predicts CLIP embeddings from fMRI data and uses diffusion-based image generation to reconstruct seen images, while Brain-Diffuser [10] combines visual rep-resentations with latent diffusion. MindEye2 further improves reconstruction through shared-subject training and predicts OpenCLIP ViT-bigG/14 representations [15, 16] for SDXL-based image generation. DREAM [17] decodes semantics, color and depth cues by reversing the pathways of the human visual system. MindTuner [12] adapts a multi-subject model to a new subject with lightweight fine-tuning, and NeuralDiffuser [13] adds guidance from primary visual features to the diffusion process. Brain-IT [6] maps brain features to image features with a transformer that uses learned queries, a design our bridges follow. Overall, these approaches demonstrate the importance of pretrained generative models for fMRI-to-image reconstruction. NeuroPictor [14] uses cortical flatmaps as input. It fine-tunes its pretrained fMRI encoder on NSD and trains a control network on its diffusion model. Our work asks whether these steps are necessary; namely, if a frozen cortical encoder paired with a frozen image generator is sufficient for competitive reconstruction.

### B. fMRI Representations

The core consideration when developing generalizable neural representations is how fMRI activity is represented. Traditional approaches reduce fMRI volumes masked with anatomical regions of interest (ROIs), producing compact feature vectors at the expense of spatial information. On the other hand, volumetric representations retain spatial structure but at a substantially greater computational cost. Recent fMRI foundation models have explored self-supervised learning to obtain general-purpose representations from large-scale datasets. For example, BrainLM [18] and BrainJEPA [19], among others, demonstrate the potential of pretrained models for downstream fMRI tasks. CortexMAE [4] frames cortical flatmaps as intermediate representations between parcellation and volumetric fMRI, converting cortical activity into 2D maps. This representation benefits from being directly compatible with Vision Transformer architectures. We use a frozen, pretrained CortexMAE encoder to turn each flatmap into a sequence of patch tokens, which our model maps to image features for reconstruction.

### C. Semantic and Low-Level Visual Features

Faithful visual reconstruction benefits from both high-level semantic information and lower-level visual structure. CLIP and OpenCLIP provide semantic visual representations that have been widely adopted for image generation and brain decoding. At the same time, convolutional feature representations such as VGG [20] retain spatial information that may be lost in semantic embeddings. Recent brain decoding approaches have consequently explored multiple visual representations. MindEye and Brain-IT use semantic representations alongside lower-level visual features to improve image reconstruction. CortexDecoder follows this general pipeline by learning separate mappings from brain representations to OpenCLIP semantic tokens and VGG features, which are subsequently combined during image generation.

## III. Cortex Decoder

CortexDecoder is a two-branch brain decoding framework that maps cortical flatmap representations to high-level semantic and low-level visual features for image reconstruction. The model first encodes cortical flatmaps using a pretrained CortexMAE encoder, producing spatially organized brain tokens. These brain tokens are then fed into two cross-attention bridges, which predict visual image features. The semantic branch predicts unpooled CLIP embeddings, while the low-level branch predicts VGG features. Finally, the predicted visual representations are provided to a frozen diffusion decoder to reconstruct the corresponding image.

Training both bridges, including the decoder-aware fine-tune, takes 3 GPU-hours on a single H100, compared with about 60 H100-hours reported for Brain-IT. All pretrained models (CortexMAE, VGG16-BN, OpenCLIP, SDXL-unCLIP) are used frozen. Both bridges are trained with AdamW (learning rate 5 *×* 10^*−*4^, weight decay 10^*−*2^). The semantic bridge is trained for 25 epochs with batch size 64 and then fine-tuned for 5 more epochs at learning rate 5 *×* 10^*−*5^ with the decoder-aware loss described in Section III-D. The low-level bridge is trained for 30 epochs with batch size 128.

### A. Natural Scenes Dataset

The Natural Scenes Dataset (NSD) is a large-scale 7T fMRI dataset in which eight participants viewed natural images drawn from MS-COCO (Microsoft Common Objects in Context) [1] across up to 40 scan sessions each. Each session comprises 750 trials. In each trial, one image is presented for 3 seconds, followed by a 1-second blank. Each subject viewed 10,000 unique images, each repeated 3 times, for a total of 30,000 trials per subject. A subset of 1000 images (shared1000) was presented to all subjects and is reserved as a standard test set throughout the literature.

The NSD experiment uses a rapid event-related design in which a new image stimulus is presented every 4 seconds, so successive stimuli occur within the timescale of the hemodynamic response. Because the responses to successive trials overlap, single-trial responses cannot be read directly from the time series. A general linear model (GLM) [21] performs this deconvolution, and its output is a static spatial map of cortical activity per trial rather than a time series. This is precisely the form that 2D flatmap representations and patch-based tokenization assume. We note that the CortexMAE encoder is spatiotemporal and retains a temporal patch dimension from its pretraining on the BOLD time series. However, because our inputs are static single-trial beta maps, we do not utilize this temporal dimension in CortexDecoder.

### B. MAE Encoder and Tokenization

NSD’s surface betas are distributed on a fsaverage mesh [22] (163,842 vertices per hemisphere). We resample them to fsLR-32k (32,492 vertices per hemisphere, 64,984 total), the standard HCP (Human Connectome Project) [23] mesh on which CortexMAE was pretrained. A pycortex-based [24] flat-surface resampler then projects these vertices onto a single-channel 224*×* 560 raster, producing the 2D cortical flatmap. Since the projection is defined by a common template mesh rather than by individual anatomy, a given location in the raster corresponds to approximately the same piece of cortex across subjects.

The flatmap is divided into non-overlapping 16*×*16 spatial patches, forming a 14 *×* 35 grid of 490 cells. Since the cortical surface occupies only part of the bounding box, 364 cells contain cortex and the remaining cells are background. CortexMAE uses a validity mask to exclude these background patches, so only 364 cortical patches are embedded.

CortexMAE is a spatiotemporal masked autoencoder [25] pretrained on BOLD time series. In other words, its patch embedding operates over both space and time. In our case, the input is a single-trial beta map, which has no temporal component. To use the pretrained model, we represent each beta map as a static 16-frame clip. Over the whole flatmap, the encoder outputs 1456 tokens of dimension 768 (364 patches *×* 4 temporal chunks). We restrict it to visual cortex by making only 50 patches that lie mostly inside NSD’s nsdgeneral mask visible to the encoder, the same way CortexMAE drops masked patches during pretraining. This gives 200 brain tokens per trial.

### C. Cross-Attention Bridge

The bridge maps the sequence of brain tokens to a sequence of visual feature tokens. Like DETR [5], the bridge uses a set of learned query vectors, with one query per output token. These are randomly initialized and trained jointly with the rest of the model. Each query is responsible for predicting one token of the target visual representation.

The brain tokens from CortexMAE serve as memory and are layer normalized to stabilize training. The queries are then passed through a stack of transformer decoder layers. Each layer applies self-attention among the queries, then cross-attention from the queries into the brain memory, and finally a position-wise feed-forward network (FFN). Self-attention lets the queries coordinate with one another so that neighboring output tokens can share information. Each query attends over all 200 brain tokens and pools the ones that carry information about its target feature. A final linear layer projects each decoded query to the dimensionality of the target representation.

We use 4 decoder layers and 8 attention heads, the optimal configuration from our size sweep (see Table 5 for other sizes). The model width is 768, with a feed-forward dimension of 3072 and dropout of 0.1. The semantic and low-level branches share these settings.

The key property of this design is that the brain representation remains a sequence throughout the pipeline. Each query attends over the full set of brain tokens. A pooled brain vector would make every query attend to the same input and reduce the decoder to a multilayer perceptron (MLP) applied identically at each output position.

We instantiate the bridge twice with one instance per branch and train the two independently. The semantic branch uses 256 queries and projects to 1664 dimensions. This matches the shape of the unpooled OpenCLIP ViT-bigG/14 [7, 15, 16] patch tokens that the diffusion decoder expects as conditioning. The low-level branch uses 7190 queries and projects to 512 dimensions. The two branches therefore differ by more than an order of magnitude in output length. They also optimize different losses, since the semantic target is dense and continuous, while the low-level target is better matched by a contrastive objective.

The semantic branch is trained with the mean squared error between the predicted and target token sequences,

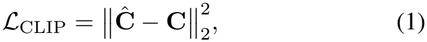

where **Ĉ**, **C** ∈ ℝ^256*×*1664^ are the predicted and target Open-CLIP tokens, and the squared error is averaged over all tokens, dimensions and trials.

To train the low-level branch, we extract target tokens from five intermediate layers of a frozen VGG16-BN network (relu1_2 through relu5_3, totaling 7,190 spatial tokens of dimension *D* = 512). For memory efficiency, at each step we randomly sample {512, 512, 128, 64, 16} spatial positions from the five layers.

At each sampled position, we use the InfoNCE loss [26] as was used in CLIP [7]. It trains the prediction for each trial to be closer to its own target than to the targets of the other trials in the batch,

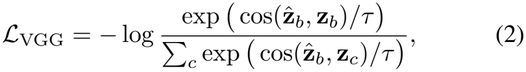

where 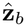 and **z**_*b*_ are the predicted and target VGG features of trial *b* at that position, *c* indexes the trials in the batch, and *τ* = 0.07. As in CLIP, we average this with the same loss with predictions and targets swapped. Because NSD repeats images, a batch can contain two trials of the same image. We drop such pairs from the denominator, since they are not true negatives. The loss is averaged over trials, sampled positions and the five layers.

### D. Semantic Branch

The semantic branch predicts the image representations that condition the diffusion decoder. Its target is the unpooled patch-token sequence produced by the OpenCLIP ViT-bigG/14 vision encoder pretrained on LAION-2B [15]. For each stimulus image, we take the 256 patch tokens before the final projection and discard the class (CLS) token. Each token is 1664-dimensional. The bridge therefore uses 256 queries and projects to 1664 dimensions so that its output matches the target.

We use an unpooled token sequence rather than a single pooled embedding. A pooled CLIP vector summarizes an image but discards its spatial layout. A token sequence retains that layout and gives the diffusion decoder a spatially structured conditioning signal. The diffusion decoder we use is a pretrained SDXL-unCLIP model whose cross-attention was trained to condition on exactly 256 bigG tokens of 1664 dimensions. Matching this shape means the decoder can be used frozen with no fine-tuning on our part. Predicting a smaller CLIP variant would require retraining the decoder’s conditioning pathway.

The OpenCLIP encoder is frozen and used only to produce targets. We extract the token sequence for every stimulus image once and cache it. During training, the branch never runs the image encoder.

MSE weighs every token dimension equally, but the frozen decoder is more sensitive to some directions of its conditioning than to others. After training with MSE alone, we therefore fine-tune the semantic bridge for 5 more epochs with an added decoder-aware term. This term is the frozen SDXL-unCLIP’s own denoising loss on the stimulus image, conditioned on the predicted tokens. It tells the bridge which predictions let the decoder reconstruct the true image. The term is weighted by *λ* = 16 (Table 4) and uses only the noise levels that our image-to-image sampling visits. The decoder stays frozen, and only the bridge is updated.

**TABLE 1.** Trained parameters of each method’s Subject 1 model, excluding frozen pretrained components. Counts are taken from released weights or from each paper.

| **Input** | **Method** | **Params** |
|---|---|---|
| Voxels | Brain-Diffuser [10] (2023) | 5.47B |
|  | MindEye1 [11] (2023) | 1.20B |
|  | MindEye2 [8] (2024) | 2.23B |
|  | MindTuner [12] (2024) | 2.24B |
|  | NeuralDiffuser [13] (2025) | 2.97B |
| Flatmap | NeuroPictor [14] (2024) | 0.676B |
|  | **CortexDecoder (ours)** | **0.083B** |

**TABLE 2.**
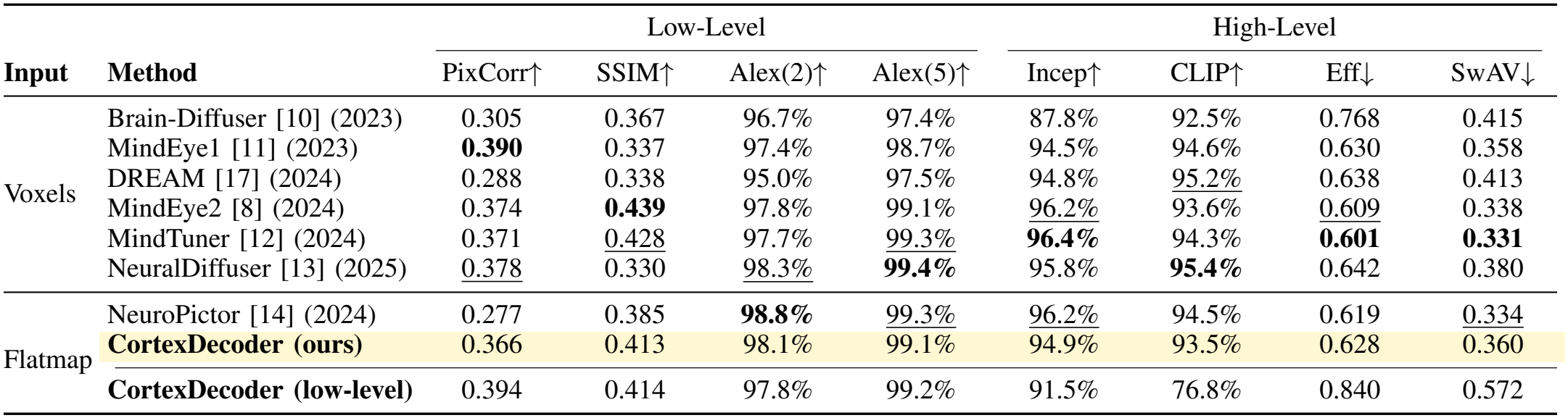
Quantitative comparison on NSD Subject 1 with baseline results taken from previous state-of-the-art methods, where MindEye2, MindTuner, NeuralDiffuser and NeuroPictor pretrain on multiple subjects before fine-tuning on Subject 1. The last row is our low-level reconstruction before diffusion (not ranked). Bold indicates the best result and underline the second best.

|  |  | Low-Level |  |  |  | High-Level |  |  |  |
|---|---|---|---|---|---|---|---|---|---|
| **Input** | **Method** | PixCorr↑ | SSIM↑ | Alex(2)↑ | Alex(5)↑ | Incep↑ | CLIP↑ | Eff↓ | SwAV↓ |
| Voxels | Brain-Diffuser [10] (2023) | 0.305 | 0.367 | 96.7% | 97.4% | 87.8% | 92.5% | 0.768 | 0.415 |
|  | MindEye1 [11] (2023) | **0.390** | 0.337 | 97.4% | 98.7% | 94.5% | 94.6% | 0.630 | 0.358 |
|  | DREAM [17] (2024) | 0.288 | 0.338 | 95.0% | 97.5% | 94.8% | <u>95.2%</u> | 0.638 | 0.413 |
|  | MindEye2 [8] (2024) | 0.374 | **0.439** | 97.8% | 99.1% | <u>96.2%</u> | 93.6% | <u>0.609</u> | 0.338 |
|  | MindTuner [12] (2024) | 0.371 | <u>0.428</u> | 97.7% | <u>99.3%</u> | **96.4%** | 94.3% | **0.601** | **0.331** |
|  | NeuralDiffuser [13] (2025) | <u>0.378</u> | 0.330 | <u>98.3%</u> | **99.4%** | 95.8% | **95.4%** | 0.642 | 0.380 |
| Flatmap | NeuroPictor [14] (2024) | 0.277 | 0.385 | **98.8%** | <u>99.3%</u> | <u>96.2%</u> | 94.5% | 0.619 | <u>0.334</u> |
|  | **CortexDecoder (ours)** | 0.366 | 0.413 | 98.1% | 99.1% | 94.9% | 93.5% | 0.628 | 0.360 |
|  | **CortexDecoder (low-level)** | 0.394 | 0.414 | 97.8% | 99.2% | 91.5% | 76.8% | 0.840 | 0.572 |

**TABLE 3.** Matched comparison on NSD Subject 1 with NeuroPictor’s frozen-encoder ablation [14]. Both methods train on Subject 1 only, keep the pretrained fMRI encoder frozen and generate one sample per image. NeuroPictor is evaluated on 982 test images and CortexDecoder on 998. Bold indicates best performance.

|  | Low-Level |  |  |  | High-Level |  |  |  |
|---|---|---|---|---|---|---|---|---|
| **Method** | PixCorr↑ | SSIM↑ | Alex(2)↑ | Alex(5)↑ | Incep↑ | CLIP↑ | Eff↓ | SwAV↓ |
| NeuroPictor (2024), frozen encoder | 0.220 | 0.342 | 97.5% | 98.8% | 93.8% | 92.8% | 0.657 | **0.360** |
| **CortexDecoder (ours)** | **0.366** | **0.413** | **98.1%** | **99.1%** | **94.9%** | **93.5%** | **0.628** | **0.360** |

**TABLE 4.** Weight *λ* of the decoder-aware fine-tuning loss. Final reconstructions on the NSD Subject 1 test set with the whole-cortex bridges and the best of 10 diffusion samples. Bold indicates the best result.

|  | Low-Level |  |  |  | High-Level |  |  |  |
|---|---|---|---|---|---|---|---|---|
| **λ** | PixCorr↑ | SSIM↑ | Alex(2)↑ | Alex(5)↑ | Incep↑ | CLIP↑ | Eff↓ | SwAV↓ |
| 0 (MSE only) | 0.372 | 0.415 | 96.7% | 97.6% | 92.8% | 91.3% | 0.673 | 0.381 |
| 4 | 0.376 | 0.416 | 97.8% | 98.6% | 93.4% | 91.9% | 0.654 | 0.374 |
| **16 (ours)** | 0.377 | 0.417 | 97.9% | **99.0%** | **94.9%** | **93.2%** | **0.640** | **0.372** |
| 32 | 0.379 | **0.422** | 98.1% | **99.0%** | 94.6% | 93.0% | 0.647 | 0.391 |
| 64 | **0.380** | 0.421 | **98.3%** | **99.0%** | 93.8% | 93.1% | 0.645 | 0.384 |

**TABLE 5.** Semantic bridge size. Token-level results are on the test set, and validation cosine is taken at the best epoch. Our configuration is in bold.

| Blocks | Heads | Params | Val. cos | Two-way |
|---|---|---|---|---|
| 2 | 8 | 20.4M | 0.523 | 99.0% |
| **4** | **8** | **39.3M** | **0.527** | **99.6%** |
| 4 | 16 | 39.3M | **0.527** | 99.4% |
| 8 | 8 | 77.1M | 0.495 | 93.2% |
| 12 | 8 | 114.9M | 0.495 | 92.8% |

**TABLE 6.** Whole-cortex input (1456 tokens) vs. visual-cortex ROI input (200 tokens) for each bridge. The DIP image depends only on the VGG bridge. All rows use *λ* = 16 and the best of 10 diffusion samples on the NSD Subject 1 test set.

|  |  | Low-Level |  |  |  | High-Level |  |  |  |
|---|---|---|---|---|---|---|---|---|---|
| **Output** | **Bridge input** | PixCorr↑ | SSIM↑ | Alex(2)↑ | Alex(5)↑ | Incep↑ | CLIP↑ | Eff↓ | SwAV↓ |
| DIP image | Whole cortex | 0.390 | 0.407 | 97.6% | 99.2% | 90.9% | 75.9% | 0.845 | 0.580 |
|  | ROI | 0.394 | 0.414 | 97.8% | 99.2% | 91.5% | 76.8% | 0.840 | 0.572 |
| Final image | Whole cortex (both bridges) | 0.377 | 0.417 | 97.9% | 99.0% | 94.9% | 93.2% | 0.640 | 0.372 |
|  | ROI semantic, whole-cortex VGG | 0.376 | 0.413 | 98.4% | 99.1% | 94.8% | 93.3% | 0.630 | 0.363 |
|  | ROI (both bridges, ours) | 0.378 | 0.417 | 98.2% | 99.0% | 94.4% | 92.9% | 0.631 | 0.363 |

### E. Low-Level Branch

From the predicted VGG feature representation, we reconstruct a low-level image through Deep Image Prior (DIP) inversion. The VGG representation consists of the stacked outputs of five convolutional layers of the VGG16-BN image model.

To convert this representation into the initial image, we optimize a separate DIP U-Net for each image. The U-Net is a small encoder-decoder convolutional generator that maps random Gaussian noise to an RGB image, which is then passed through a frozen VGG16-BN network. The extracted VGG features are compared against the brain-predicted VGG representation using an MSE loss. The gradient is backpropagated only through the DIP U-Net, iteratively updating the generator for a fixed 2000 optimization steps to produce the final low-level reconstruction.

### F. Final Image Reconstruction

To generate the final image reconstruction, we combine the image capturing low-level features generated using DIP with the semantic CLIP prediction using the MindEye2 Stable Diffusion XL unCLIP model (SDXL-unCLIP). SDXL-unCLIP is a pretrained image generation model that generates images from OpenCLIP ViT-bigG/14 image embeddings instead of text-prompt embeddings. The intention is for the low-level DIP image to provide layout, color, and coarse structure while the predicted OpenCLIP ViT-bigG/14 tokens contribute the semantic context of the image.

We first add Gaussian noise to the low-level image to partially corrupt fine detail while preserving coarse layout and color. This re-noised image becomes the initial state for SDXL-unCLIP, anchoring the image structure. The predicted OpenCLIP token representation then steers the denoising process toward the correct semantic context, turning the blurry low-level reconstruction into a detailed final image. With this approach, the reconstruction should reflect both the content and the structure of the stimulus.

## IV. Results

We evaluate CortexDecoder on the NSD shared1000 test set (the 998 shared images that Subject 1 viewed), which is kept separate from the training and validation data to prevent leakage. The metrics compare the reconstructions against the ground truth stimulus images. All approaches detailed here were evaluated on Subject 1. We report PixCorr (Pixel Correlation), SSIM (Structural Similarity Index Measure) [27], AlexNet(2), AlexNet(5) [28], InceptionV3 [29], CLIP [7], EfficientNet-B1 [30] and SwAV-ResNet50 [31]. PixCorr and SSIM measure pixel and structural similarity while AlexNet, InceptionV3 and CLIP measure feature-space similarity. These three are reported as two-way identification accuracy, which is the percentage of comparisons in which a reconstruction’s features are closer to its own stimulus than to another test image (chance = 50%). EfficientNet and SwAV report correlation distance, where lower is better. Benchmark metrics from the subjects 2, 5, and 7 are included in the Table 7 and Table 8.

**TABLE 7.**
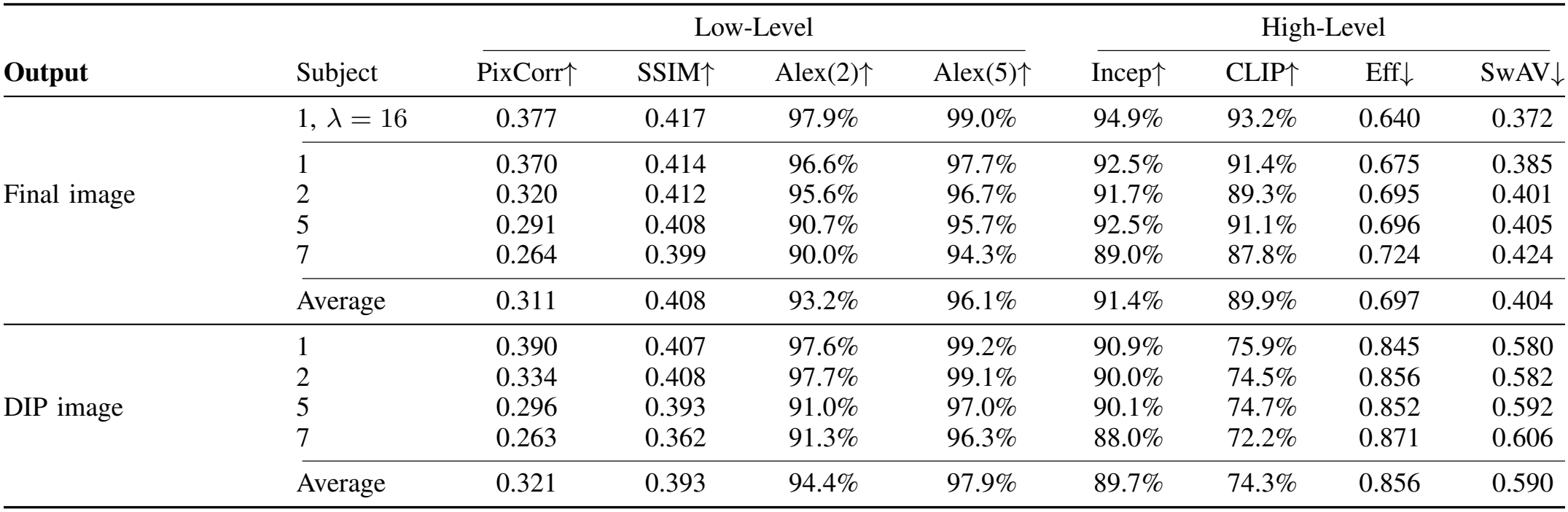
CortexDecoder on Subjects 1, 2, 5 and 7 with the whole-cortex bridges and the best of 10 diffusion samples. The row *λ* = 16 adds the decoder-aware fine-tune.

| **Output** | Subject | Low-Level |  |  |  | High-Level |  |  |  |
|---|---|---|---|---|---|---|---|---|---|
|  |  | PixCorr↑ | SSIM↑ | Alex(2)↑ | Alex(5)↑ | Incep↑ | CLIP↑ | Eff↓ | SwAV↓ |
| Final image | 1, λ = 16 | 0.377 | 0.417 | 97.9% | 99.0% | 94.9% | 93.2% | 0.640 | 0.372 |
|  | 1 | 0.370 | 0.414 | 96.6% | 97.7% | 92.5% | 91.4% | 0.675 | 0.385 |
|  | 2 | 0.320 | 0.412 | 95.6% | 96.7% | 91.7% | 89.3% | 0.695 | 0.401 |
|  | 5 | 0.291 | 0.408 | 90.7% | 95.7% | 92.5% | 91.1% | 0.696 | 0.405 |
|  | 7 | 0.264 | 0.399 | 90.0% | 94.3% | 89.0% | 87.8% | 0.724 | 0.424 |
|  | Average | 0.311 | 0.408 | 93.2% | 96.1% | 91.4% | 89.9% | 0.697 | 0.404 |
| DIP image | 1 | 0.390 | 0.407 | 97.6% | 99.2% | 90.9% | 75.9% | 0.845 | 0.580 |
|  | 2 | 0.334 | 0.408 | 97.7% | 99.1% | 90.0% | 74.5% | 0.856 | 0.582 |
|  | 5 | 0.296 | 0.393 | 91.0% | 97.0% | 90.1% | 74.7% | 0.852 | 0.592 |
|  | 7 | 0.263 | 0.362 | 91.3% | 96.3% | 88.0% | 72.2% | 0.871 | 0.606 |
|  | Average | 0.321 | 0.393 | 94.4% | 97.9% | 89.7% | 74.3% | 0.856 | 0.590 |

**TABLE 8.** Per-subject results of prior methods, taken from their papers. Averages are computed from these values and can differ from each paper’s own average by rounding.

|  |  | Low-Level |  |  |  | High-Level |  |  |  |
|---|---|---|---|---|---|---|---|---|---|
| **Method** | Subject | PixCorr↑ | SSIM↑ | Alex(2)↑ | Alex(5)↑ | Incep↑ | CLIP↑ | Eff↓ | SwAV↓ |
| MindEye2 [8] | 1 | 0.374 | 0.439 | 97.8% | 99.1% | 96.2% | 93.6% | 0.609 | 0.338 |
|  | 2 | 0.328 | 0.430 | 97.0% | 98.8% | 94.9% | 91.7% | 0.631 | 0.347 |
|  | 5 | 0.301 | 0.432 | 95.3% | 98.5% | 96.5% | 94.3% | 0.600 | 0.335 |
|  | 7 | 0.283 | 0.423 | 94.3% | 98.0% | 94.1% | 92.4% | 0.638 | 0.357 |
|  | Average | 0.322 | 0.431 | 96.1% | 98.6% | 95.4% | 93.0% | 0.620 | 0.344 |
| MindTuner [12] | 1 | 0.371 | 0.428 | 97.7% | 99.3% | 96.4% | 94.3% | 0.601 | 0.331 |
|  | 2 | 0.331 | 0.421 | 96.8% | 99.0% | 95.1% | 92.7% | 0.620 | 0.342 |
|  | 5 | 0.298 | 0.422 | 94.7% | 98.7% | 96.6% | 94.7% | 0.592 | 0.333 |
|  | 7 | 0.288 | 0.414 | 94.1% | 98.1% | 94.1% | 93.3% | 0.636 | 0.355 |
|  | Average | 0.322 | 0.421 | 95.8% | 98.8% | 95.6% | 93.8% | 0.612 | 0.340 |
| NeuralDiffuser [13] | 1 | 0.378 | 0.330 | 98.3% | 99.4% | 95.8% | 95.4% | 0.642 | 0.380 |
|  | 2 | 0.309 | 0.318 | 96.6% | 98.7% | 94.7% | 95.0% | 0.670 | 0.404 |
|  | 5 | 0.261 | 0.315 | 93.7% | 97.9% | 96.5% | 96.9% | 0.658 | 0.402 |
|  | 7 | 0.247 | 0.310 | 92.8% | 97.1% | 94.6% | 94.5% | 0.683 | 0.418 |
|  | Average | 0.299 | 0.318 | 95.3% | 98.3% | 95.4% | 95.4% | 0.663 | 0.401 |
| DREAM [17] | 1 | 0.288 | 0.338 | 95.0% | 97.5% | 94.8% | 95.2% | 0.638 | 0.413 |
|  | 2 | 0.273 | 0.331 | 94.2% | 97.1% | 93.4% | 93.5% | 0.652 | 0.422 |
|  | 5 | 0.269 | 0.325 | 93.5% | 96.6% | 93.8% | 94.1% | 0.633 | 0.397 |
|  | 7 | 0.265 | 0.319 | 92.7% | 95.4% | 92.6% | 93.7% | 0.656 | 0.438 |
|  | Average | 0.274 | 0.328 | 93.9% | 96.7% | 93.7% | 94.1% | 0.645 | 0.418 |

We report metrics for both the low-level DIP reconstruction and the final diffusion reconstruction (Fig. 5). This allows us to evaluate the contribution of the low-level branch separately from the semantic conditioning. We generate a single diffusion sample per image.

**Fig 1.**
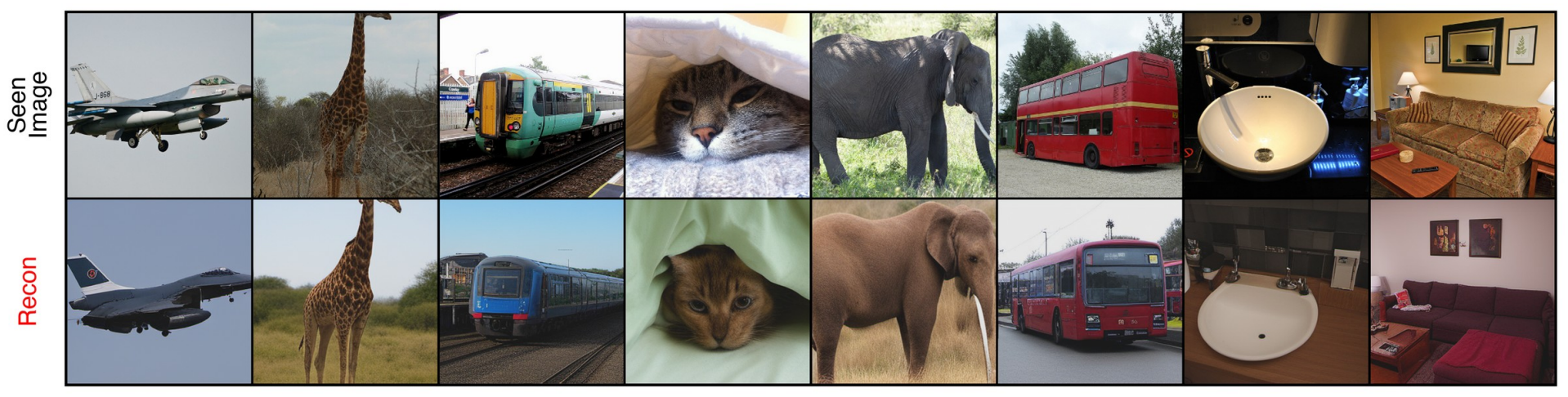
Stimuli (top) and CortexDecoder reconstructions (bottom). Images from COCO [1] (CC BY 4.0).

**Fig 2.**
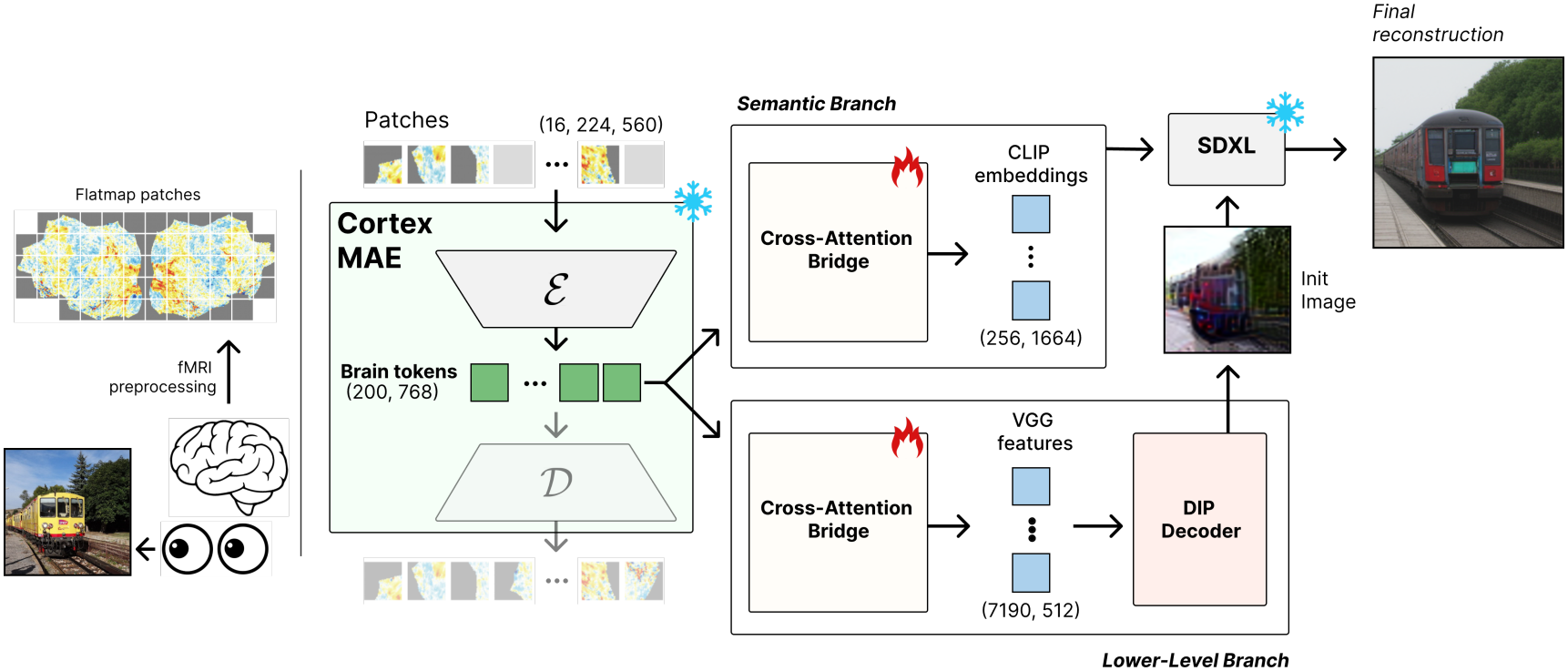
CortexDecoder overview. A frozen CortexMAE encoder turns the visual-cortex patches of the cortical flatmap into 200 brain tokens. Two cross-attention bridges, the only trained components, predict OpenCLIP tokens (semantic) and VGG16-BN features (low-level). A DIP-inverted low-level image initializes SDXL-unCLIP, which is conditioned on the predicted OpenCLIP tokens to produce the final image. Images from COCO [1] (CC BY 4.0).

**Fig 3.**
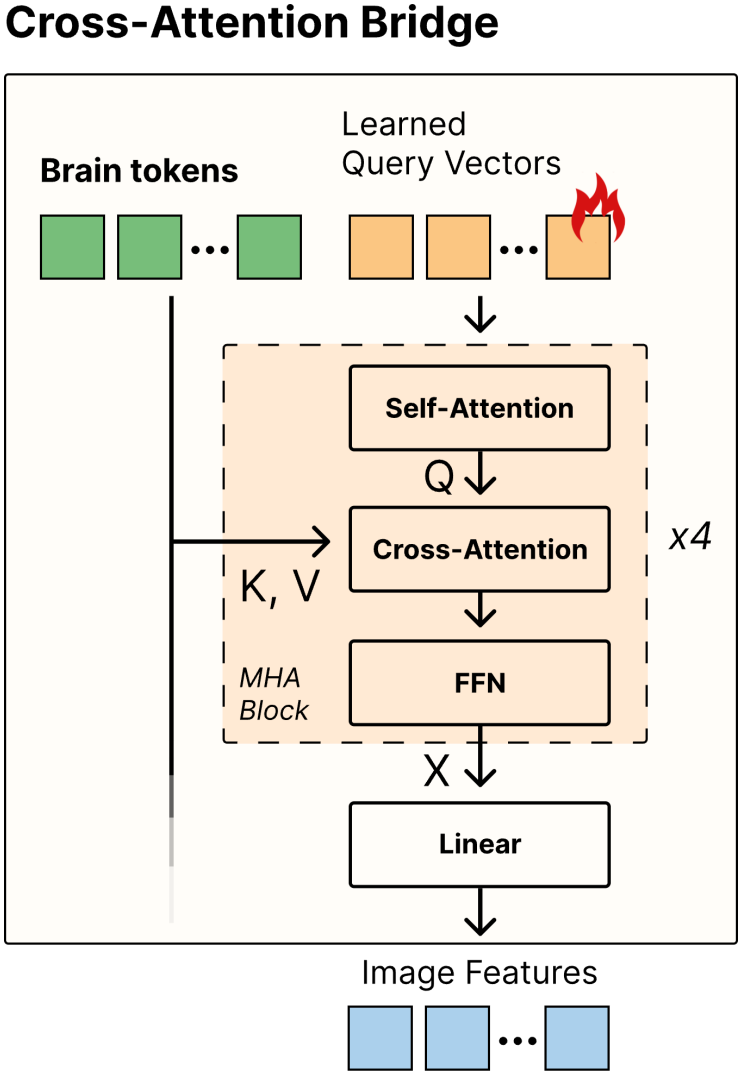
Cross-attention bridge. Learned queries, one per output token, attend to the brain tokens through 4 blocks of self-attention, cross-attention and FFN. A linear layer projects each query to the target feature dimension. Each branch has its own bridge.

**Fig 4.**
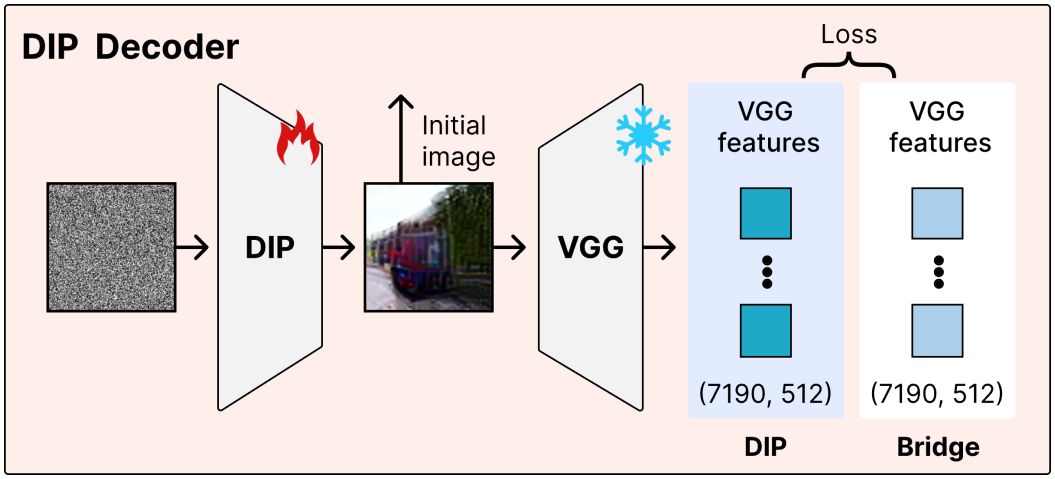
Low-level reconstruction with Deep Image Prior. A U-Net maps noise to an image whose frozen VGG16-BN features are matched to the bridge’s predicted features. After 2000 optimization steps, its output is the low-level reconstruction.

**Fig 5.**
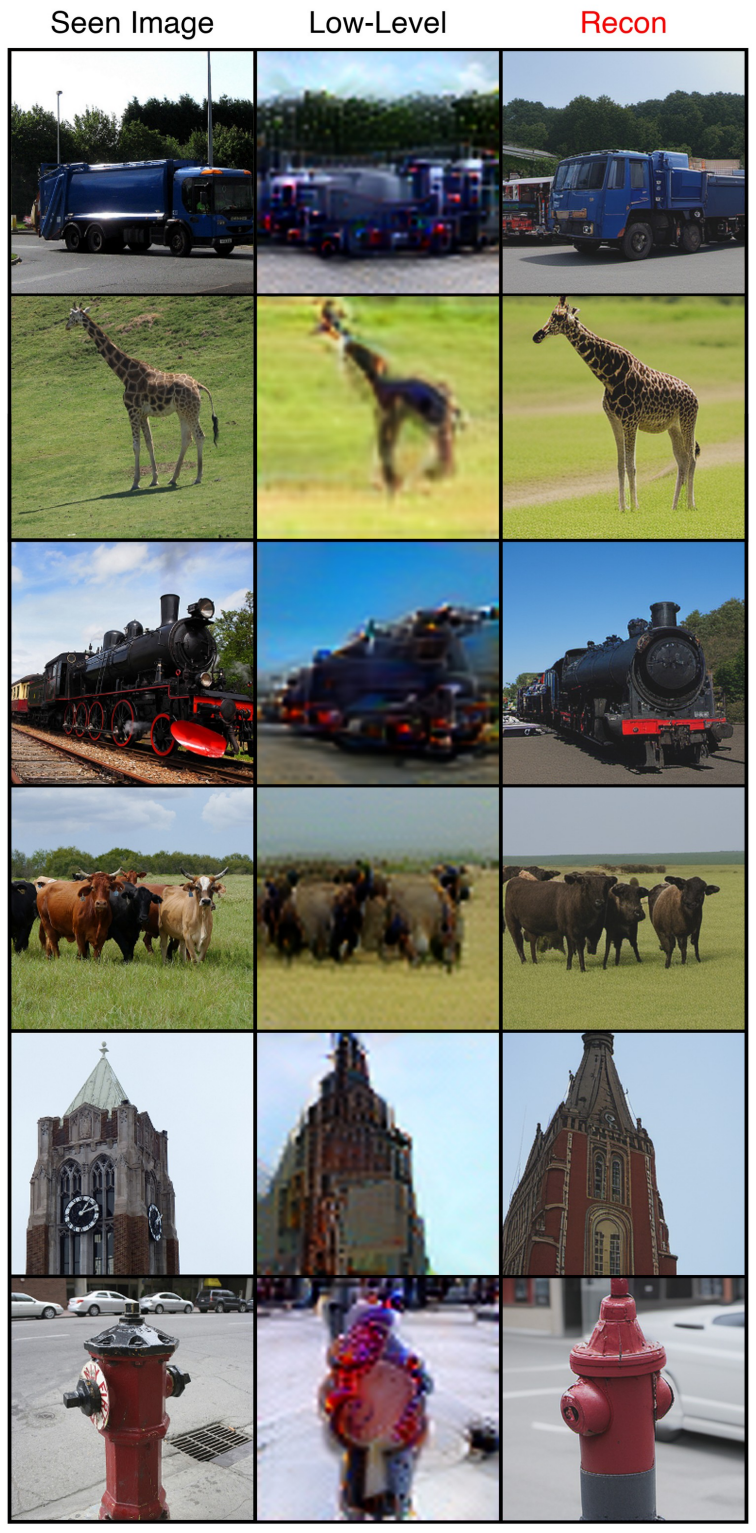
Stimulus (left), low-level DIP reconstruction (middle) and final diffusion reconstruction (right). Images from COCO [1] (CC BY 4.0).

### A. Reconstruction Benchmarks

CortexDecoder is competitive with prior methods in capturing low-level and mid-level features in reconstructed images (Table 2). Its SSIM (0.413) and AlexNet(2) and AlexNet(5) scores (98.1% and 99.1%) are close to the best methods, reflecting the contribution of the low-level branch. However, it is weaker on semantic metrics, particularly CLIP.

Four of the prior methods in Table 2 pretrain on multiple subjects, and NeuroPictor also fine-tunes its fMRI encoder. For a matched comparison, Table 3 compares CortexDecoder with NeuroPictor’s ablation that trains on Subject 1 only and keeps its pretrained fMRI encoder frozen, the same setting as ours. Under these conditions, CortexDecoder outperforms NeuroPictor on seven of the eight metrics and ties on SwAV, with the largest gains on PixCorr (0.366 vs. 0.220) and SSIM (0.413 vs. 0.342). NeuroPictor’s ablation still trains a control network on its diffusion model, while our generator is frozen. Fig. 6 shows reconstructions from CortexDecoder, MindEye2 and NeuroPictor for the same stimuli.

The results show that introducing semantic conditioning degrades low-level fidelity. The low-level reconstruction on its own scores higher on most low-level metrics (PixCorr, SSIM and AlexNet(5)) than the final reconstruction, where semantic conditioning is introduced in the diffusion stage. We conclude that the final reconstruction, which combines semantic and low-level representations, improves on semantic metrics at the expense of low-level performance.

**Fig 6.**
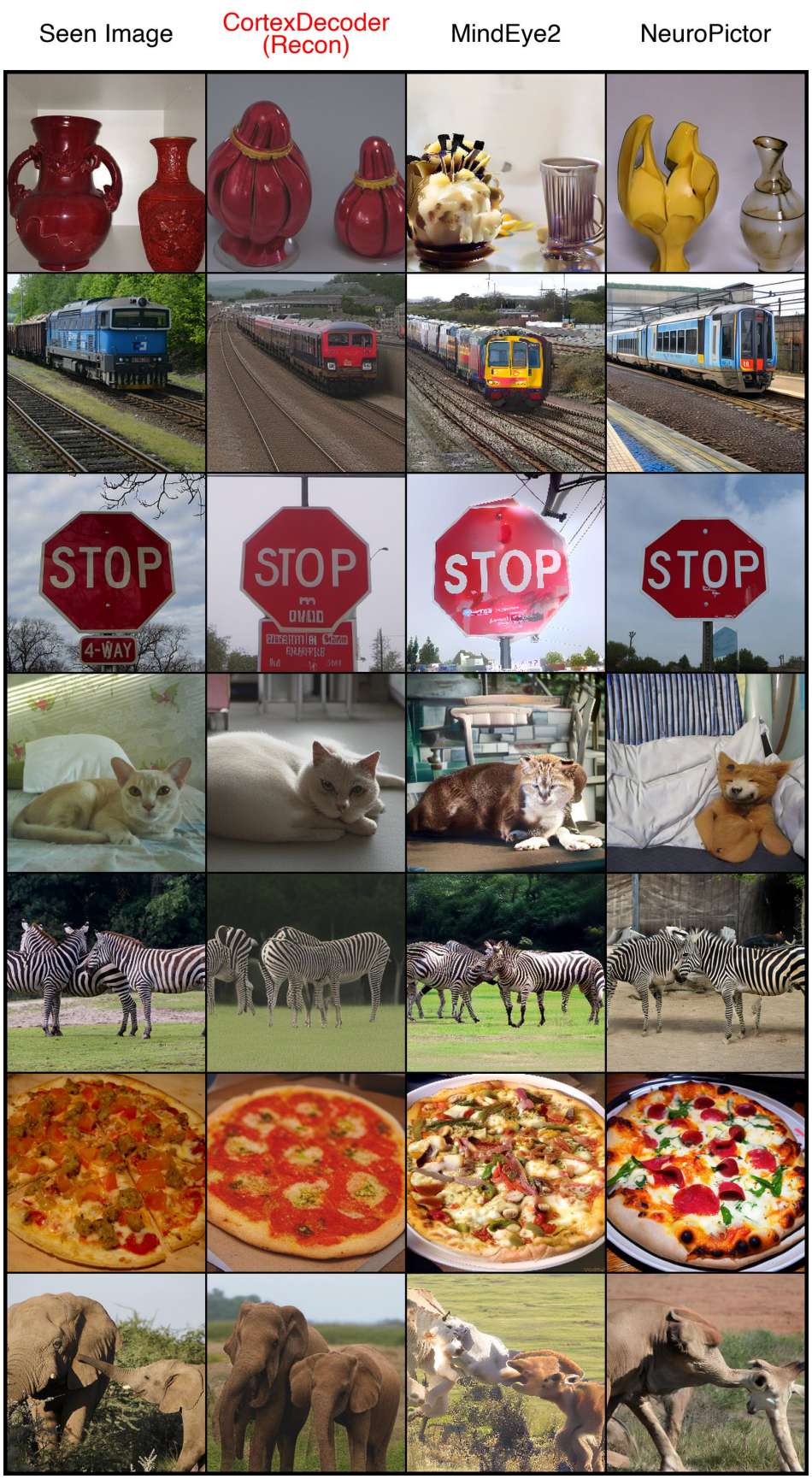
Stimulus, CortexDecoder, MindEye2 and NeuroPictor reconstructions (left to right). Images from COCO [1] (CC BY 4.0).

**Fig 7.**
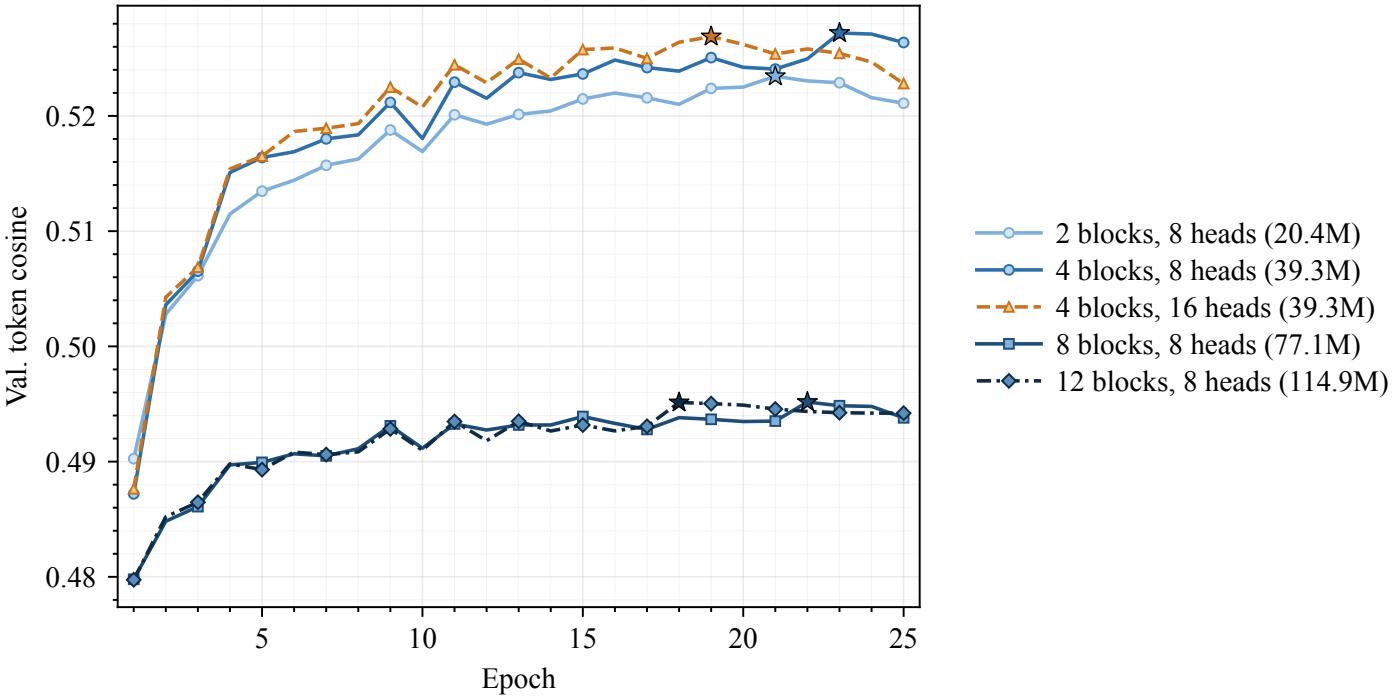
Validation token cosine per epoch for each semantic bridge size in Table 5. Stars mark the best epoch of each run, whose checkpoint is kept.

**Fig 8.**
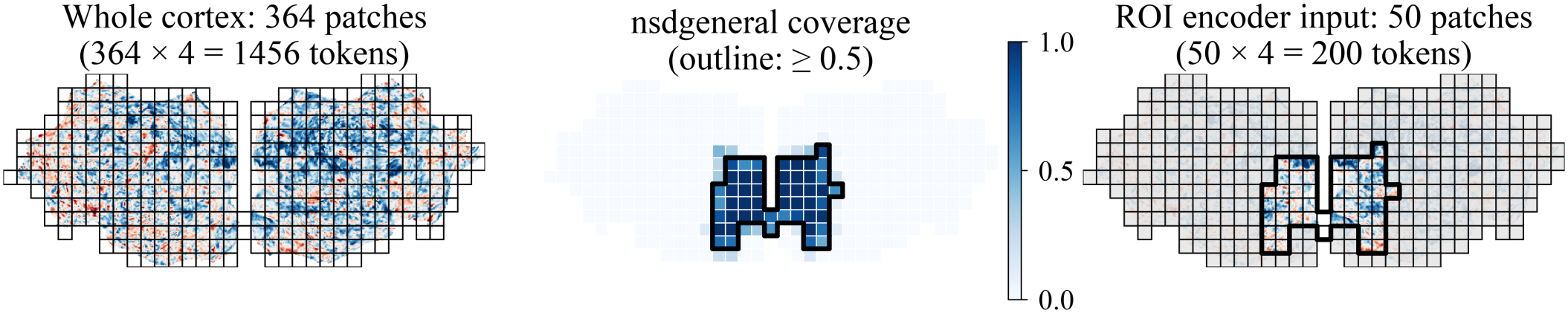
Visual cortex restriction on the CortexMAE patch grid. The left panel shows a single-trial Subject 1 beta flatmap with all 364 cortical patches. The middle panel shows the fraction of each patch inside NSD’s nsdgeneral mask. The right panel shows the same flatmap with only the 50 patches that have at least half their area inside nsdgeneral, which are the only patches visible to the encoder.

**Fig 9.**
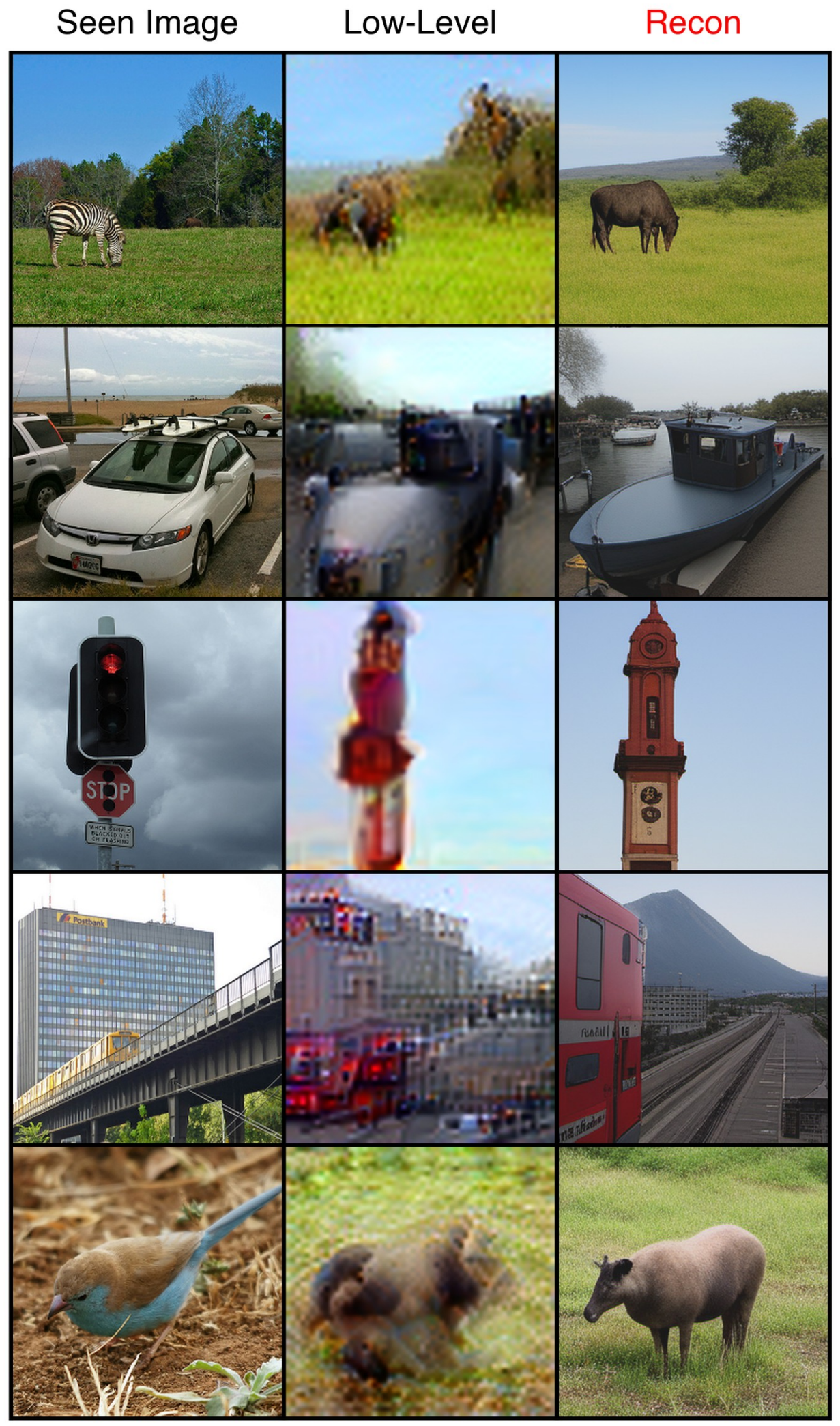
Failure cases. Stimulus (left), low-level DIP reconstruction (middle) and final reconstruction (right). In these examples, the low-level image is already wrong. When its layout or colors are off, the final image shows a plausible but wrong object. Images from COCO [1] (CC BY 4.0).

**Fig 10.**
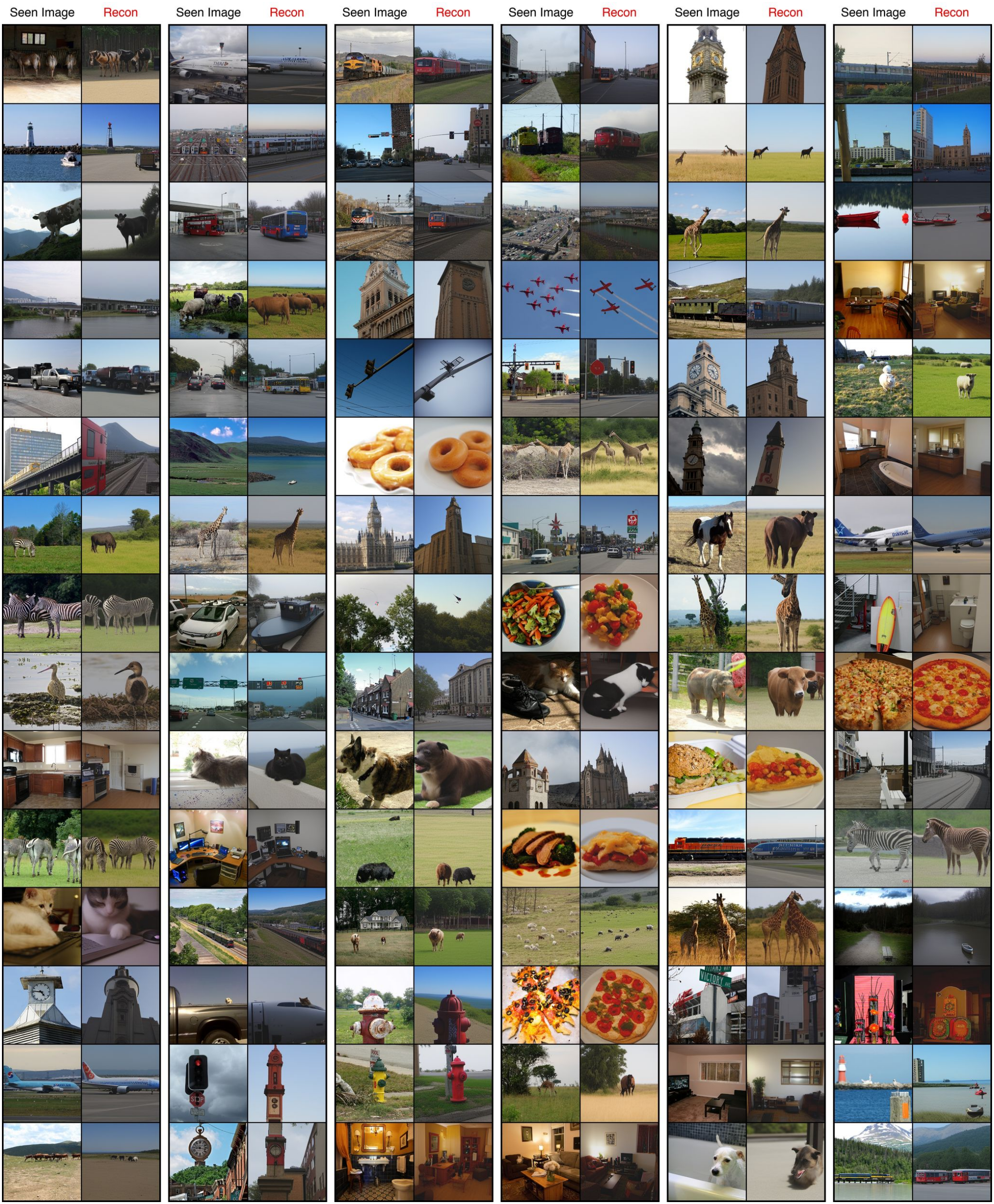
CortexDecoder reconstructions for NSD Subject 1 test images. Each pair shows the stimulus (left) and the final reconstruction (right). Images from COCO [1] (CC BY 4.0).

**Fig 11.**
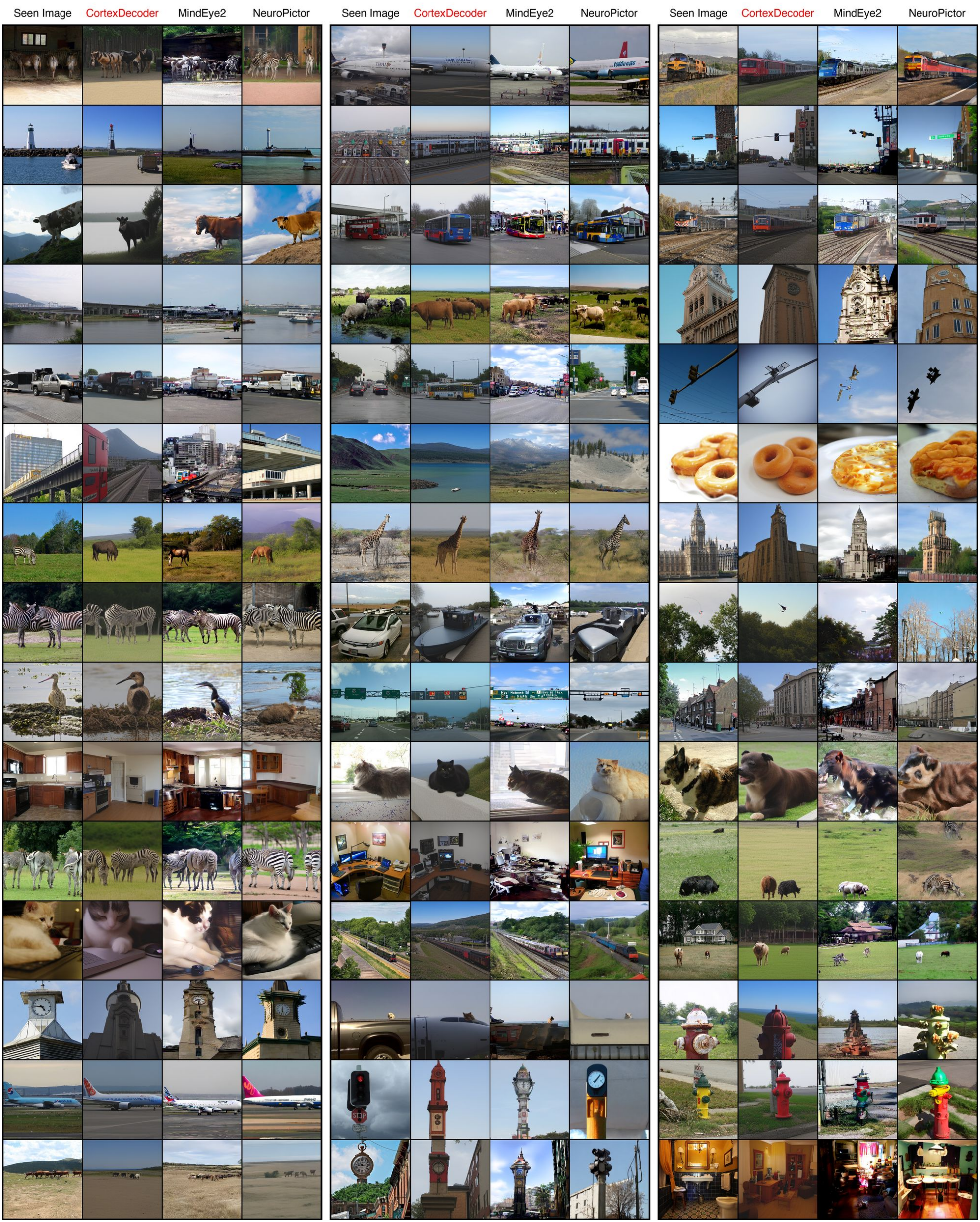
Stimulus, CortexDecoder, MindEye2 and NeuroPictor reconstructions for NSD Subject 1 test images. MindEye2 and NeuroPictor reconstructions are from their publicly released Subject 1 outputs. Images from COCO [1] (CC BY 4.0).

## V. Conclusion

We introduce CortexDecoder, an fMRI-to-image reconstruction framework that uses 2D cortical flatmaps as neural representations for visual decoding. By converting cortical flatmaps into patch tokens, CortexDecoder enables the use of Vision Transformers while preserving the spatial structure of cortical responses. A pretrained CortexMAE encoder produces brain tokens, which are then mapped through two cross-attention branches, one for semantic features and one for low-level image reconstruction. These predictions are then passed to a frozen SDXL-unCLIP image generation model, which takes the predicted low-level image as its initial image and generates the final reconstruction conditioned on the predicted semantic embedding.

Our results show that cortical flatmaps preserve enough information to support brain decoding, which corroborates NeuroPictor’s prior work [14]. We demonstrate that utilizing pretrained frozen foundation models together with two lightweight trainable bridges is enough to achieve competitive benchmarks among other decoding frameworks.

## VI. Limitations

Our work demonstrates that lightweight models designed to bridge representations, when paired with frozen foundation models like CortexMAE and SDXL, are sufficient to achieve competitive evaluations. However, several limitations remain, including occasional semantic and low-level image reconstruction errors which are shown in Fig. 9. It remains a fact that our model trains on one subject at a time and is subject specific. However, we believe CortexDecoder’s lightweight nature makes it viable to adapt new subjects in a tractable manner due to its relatively inexpensive training process. Lastly, this model expects fMRI as input which requires extensive data pre-processing limiting its practical use compared to other lightweight brain recording devices such as EEG.

## Appendix A

## Appendix B

## Appendix C

## Appendix D

### Data and Code Availability

Our dataset is available at https://huggingface.co/datasets/teooi/cortexdecoder-nsd-flatmaps. Code is available at https://github.com/teooi/CortexDecoder. All experiments were run on Modal cloud instances with a single NVIDIA H100 (80 GB) GPU and 64 GB of system RAM, running Debian Linux with Python 3.11, PyTorch 2.8.0 and CUDA 12.9. All random seeds were set to 42.

